# Peptide Scanning of SARS-CoV and SARS-CoV-2 Spike Protein Subunit 1 Reveals Potential Additional Receptor Binding Sites

**DOI:** 10.1101/2021.08.16.456470

**Authors:** Weilin Lin, Jannatul Rafeya, Vanessa Roschewitz, David Smith, Adrian Keller, Yixin Zhang

## Abstract

The binding of SARS-CoV and SARS-CoV-2 to the ACE2 receptor on human cells is mediated by the spike protein subunit 1 (S1) on the virus surfaces, while the receptor binding domains (RBDs) of S1 are the major determinants for the interaction with ACE2 and dominant targets of neutralizing antibodies. However, at the virus-host interface, additional biomolecular interactions, although being relatively weak in affinity and low in specificity, could also contribute to viral attachment and play important roles in gain- or loss-of-function mutations. In this work, we performed a peptide scanning of the S1 domains of SARS-CoV and SARS-CoV-2 by synthesizing 972 16-mer native and mutated peptide fragments using a high throughput *in situ* array synthesis technology. By probing the array using fluorescently labelled ACE2, a number of previously unknown potential receptor binding sites of S1 have been revealed. 20 peptides were synthesized using solid phase peptide synthesis, in order to validate and quantify their binding to ACE2. Four ACE2-binding peptides were selected, to investigate whether they can be assembled through a biotinylated peptide/neutravidin system to achieve high affinity to ACE2. A number of constructs exhibited high affinity to ACE2 with K_d_ values of pM to low nM.

## Introduction

The COVID-19 pandemic, caused by infection with the severe acute respiratory syndrome related coronavirus-2 (SARS-CoV-2), has led to millions of fatalities as well as devastating social and economic consequences[1]. Molecular and structural biology studies have revealed the most potent and specific interaction between the virus and human cells, namely, the interaction between the RBD domain of viral spike protein subunit 1 (S1) and the receptor ACE2 (angiotensin converting enzyme 2) on human cells[2–4]. Despite its important function, the RBD is highly variable among sarbecoviruses, as compared to many other sequences of viral proteins[2]. SARS-CoV-2 is 82% similar to the SARS-CoV attributed to the SARS outbreak in 2003, while their similarities of the RBD domains and receptor binding motifs (RBM), i.e., the 69-residue motif in the RBD directly interacting with the receptor, are 67% and 47%, respectively. Other viral proteins, as well as parts of the S1 other than the RBD can also contribute to the interaction, while other human proteins have also been found to mediate the viral entry. For example, a monoclonal antibody against the N-terminal domain of S1 (NTD) has been found to able to block the infection effectively, though it does not inhibit the binding of S1 to its receptor on cells[5]. Human serine protease TMPRSS2 is used for S protein priming and TMPRSS2 inhibitor can block the viral entry[6].

Comprehensive profiling of the interactions between the viral proteins and the proteins on human cells is essential in understanding the infection mechanism and to develop diagnostic tests and therapies. There is no single technology, which will allow us to elucidate all these interactions at molecular detail, especially regarding the transmission and evolution of viruses with host cells from different species. Proteomic and genetic studies were used to map the potential interactions[7, 8], while crystallography and cryo-EM can help to illustrate the individual interactions with atomic to sub-nm resolution[3, 9]. As a chemical biology approach, peptide arrays have the advantage to map the sequence-specific interactions quickly, though the peptide fragments often cannot fully resemble the structures of folded proteins. Moreover, the identified peptides can become lead compounds for the development of antiviral drugs to block the cellular entry of viruses, a therapeutic approach complementary to vaccines[10]. Peptide arrays have also been used to map the viral epitopes against antivirus antibodies[11].

The coronavirus spike proteins are homotrimeric class I viral fusion proteins containing S1 and S2 subunits. For membrane fusion, the spike proteins need to be proteolytically activated at the S1/S2 boundary, resulting in the dissociation of S1 and a dramatic structural change of S2. Different from SARS-CoV, cell entry of SARS-CoV-2 can be pre-activated by proprotein convertase furin, reducing its dependence on target cell proteases for entry. The interaction between S1 and ACE2 is essential for initial virus attachment. S1 can be divided into an N-terminal domain (NTD) and a C-terminal domain (CTD), while the CTD can be further divided into RBD and two subdomains SD1 and SD2. The RBD is conformationally variable. In the closed conformation, the RBDs are shielded by the NTDs. In the open conformation, one RBD is exposed upward away from the viral membrane. The SARS-CoV-2 RBD has a twisted five-stranded antiparallel β sheet. Between the β4 and β7 strands, there is an extended insertion, known as receptor binding motif (RBM), containing the short β5 and β6 strands, α4 and α5 helices, and loops. Multiple residues of the SARS-CoV-2 RBM have been found essential for contacting ACE2.

The structure of spike protein is considered to be highly dynamic, not only the RBD, but also the other subunits and domains, allowing it to search for the receptor for docking to the target cell. Therefore, interactions other than those observed in the final complex can be involved in the initial binding of the viruses to host cells. To identify these potential interactions by scanning the S1 domains of SARS-CoV and SARS-CoV-2, in this work, we prepared peptide arrays representing the two S1 proteins, in which each peptide was 16 amino acids long with a 2 amino acids offset and 14 amino acids overlap (figure 1A). All reported mutations prior to May 2021 have also been included in this study. The resulting arrays were then probed by fluorescently labelled ACE2.

**Figure 1.**
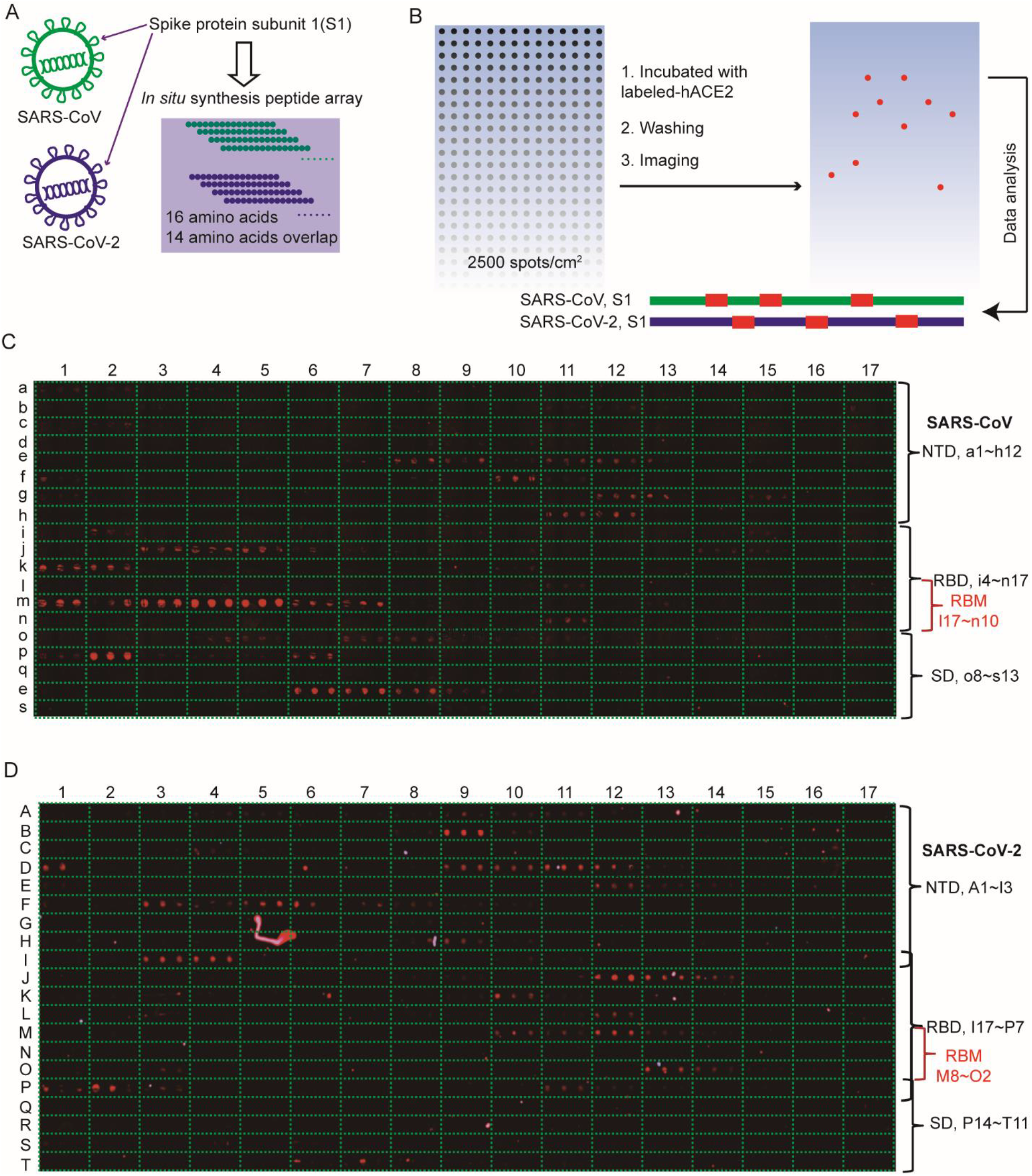
SARS-CoV and SARS-CoV-2 spike protein S1 subunit microarrays for the mapping of ACE2 interactions. (A, B) The schematic illustration of mapping peptides from S1 subunits and ACE2 interactions. (C,D) Binding paterns of peptides from SARS-CoV (C) and SARS-CoV-2 (D) S1 subunits to ACE2.

## Results and discussion

We have recently developed an amphiphilic hydrogel coating for glass surfaces, on which small droplets of polar aprotic organic solvents can be deposited with relatively large contact angle (> 40°) and inhibited motion, permitting multiple rounds of on-demand combinatorial synthesis of small molecule compounds and peptides with high yields [12]. Moreover, the amphiphilic hydrogel coating can be switched to a hydrophilic matrix after the synthesis, resulting in a hydrogel coating suitable for protein- and cell-based screening. Automation of the array synthesis allowed us to generate high density peptide arrays with standard solid phase peptide chemistry. 334 peptides, each in triplicate, presenting the amino acids 14-695 of SARS-CoV-2 S1, and 319 peptides presenting the amino acids 18-669 of SARS-CoV S1 were synthesized on the hydrogel coating (figure 1A). The resulting arrays possess a density of 2500 features/cm^2^. Because of the small size of the chip (about 1 cm^2^), very small amount of protein was needed for the screening (200 ng of fluorescently labelled human ACE2 in 5 μL). After washing with buffer to remove the unbound protein, the glass slide was imaged using afluorescence microscope (figure 1B).

As shown in figure 1C and 1D, the hydrogel coating does not show any non-specific interactions with ACE2. Seven successive peptide fragments (m1-m7, figure 1C) from the RBM of SARS-CoV showed strong signals in binding to ACE2, while the strongest signals were observed for the peptides m3-m5 (figure 1C). Other binding sites of SARS-CoV S1 to ACE2, both in the RBD and in the other domains, have also been identified. Interestingly, for the SARS-CoV-2 RBM peptides, only three successive peptide fragments have shown binding signals (M10-M12, figure 1D), while the strongest signals were from outside the RBM, e.g. the peptide J12 (figure 1D) of the RBD and the peptide B9 (figure 1D) of the NTD. After sequence alignment, we found that both viruses share a similar binding profile to ACE2 in the RBD sequences, but exhibit remarkable differences in the other domains (see figure S1).

We selected 21 peptides to measure their binding affinities to ACE2 (figure 2). The peptides were synthesized using standard solid phase peptide chemistry and purified by HPLC. The binding affinities were measured using bio-layer interferometry with ACE2 immobilized on the sensor. As the sequences were identified with the peptides on a solid support and the proteins in solution, validating their interaction with proteins immobilized on sensor surface and the peptides in the mobile phase can avoid artifacts caused by multi-valent interactions[13]. Four SARS-CoV-2 peptides and three SARS-CoV peptides showed strong interaction with ACE2. Interestingly, only the NTD peptides from SARS-CoV-2, but not the NTD peptides from SARS-CoV, exhibited strong interaction with the receptor protein. The strongest interaction was measured not with the RBM peptide of SARS-CoV-2, but with an RBD peptide of SARS-CoV-2.

**Figure 2.**
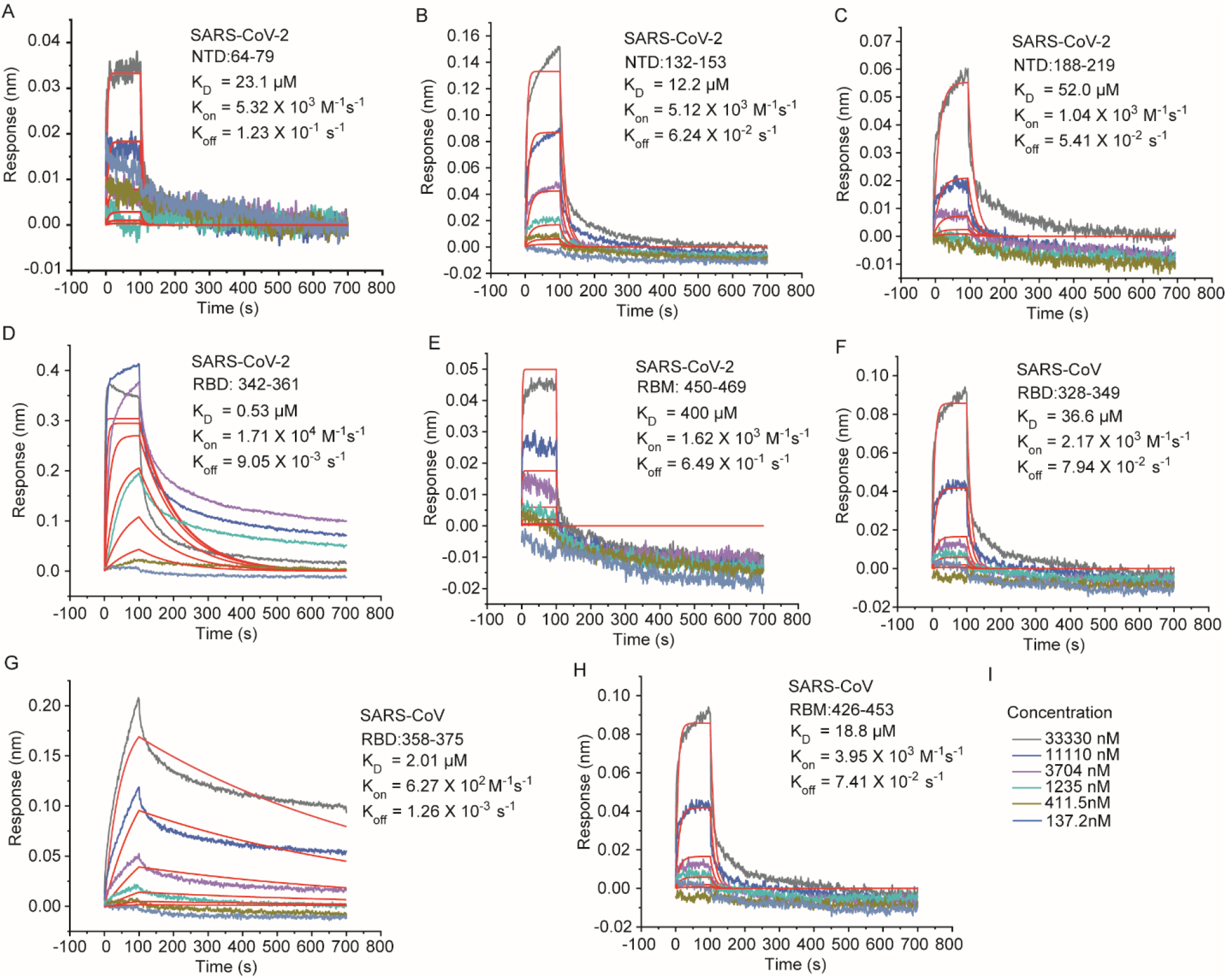
Binding kinetics of peptides from SARS-CoV (A-E) and SARS-CoV-2 (F-H) S1 subunits to ACE2. ACE2 was immobilized on BLI biosensors and measured for binding with various concentrations of the peptides.

As the peptide fragments cannot fully resemble their structures in the folded proteins, the patterns of fluorescence signals on the arrays and the measured binding affinities do not contradict the interaction between RBM and ACE2 observed in the crystal structures. Meanwhile, peptide scanning can help to reveal the relatively weak interactions, which cannot be seen in the final complex structure. The structure of the spike proteins is highly dynamic, and mechanisms associated with conformational changes have been suggested to contribute to viral attachment and cell entry[14]. The interaction between the non-RBM peptides with ACE2 can be transient and relatively weak, but relevant to the pathology of the viruses. Both array screening and affinity measurements have shown different patterns between the SARS-CoV and SARS-CoV-2 peptides. This might reflect their difference in infection, while the dynamic processes cannot be easily visualized by conventional structural biology methods. We then tested the effects of mutations on array screening. The sequences of SARS-CoV-2 S1 mutants were extracted from the CDC database[15]. The 13 strains include most mutations prior to May 12, 2021. All peptide sequences containing the mutations have been synthesized, resulting in a secondary array of 319 different peptides (each in triplicate). Figure 3 summarizes all mutations, which can cause changes in binding to ACE2. Nine mutations have been found to cause an increase of ACE2 binding in one or multiple peptide fragments in the scan (figure 3A). Six mutations have shown an effect of reduced ACE2 binding in one or multiple peptide fragments (figure 3B). Both increase and decrease of ACE2 binding were observed in the peptide scanning of four mutations (figure 3C). It is important to note that many changes, especially those causing increased binding, have been measured for multiple successive sequences (e.g. P26S, Δ156, E154K, L452R, and E484I), a common indication of reliability of peptide scanning. Among the multiple mutations, remarkable changes in the peptide scanning patterns have been observed for A67V+Δ69/70 and S474N+S484K (figure 3d). As most of these mutations are not in the RBD and only one mutation is in the RBM, the evolution of SARS-CoV-2 is not necessarily in the direction of increasing the binding affinity between S1 and its receptor.

**Figure 3.**
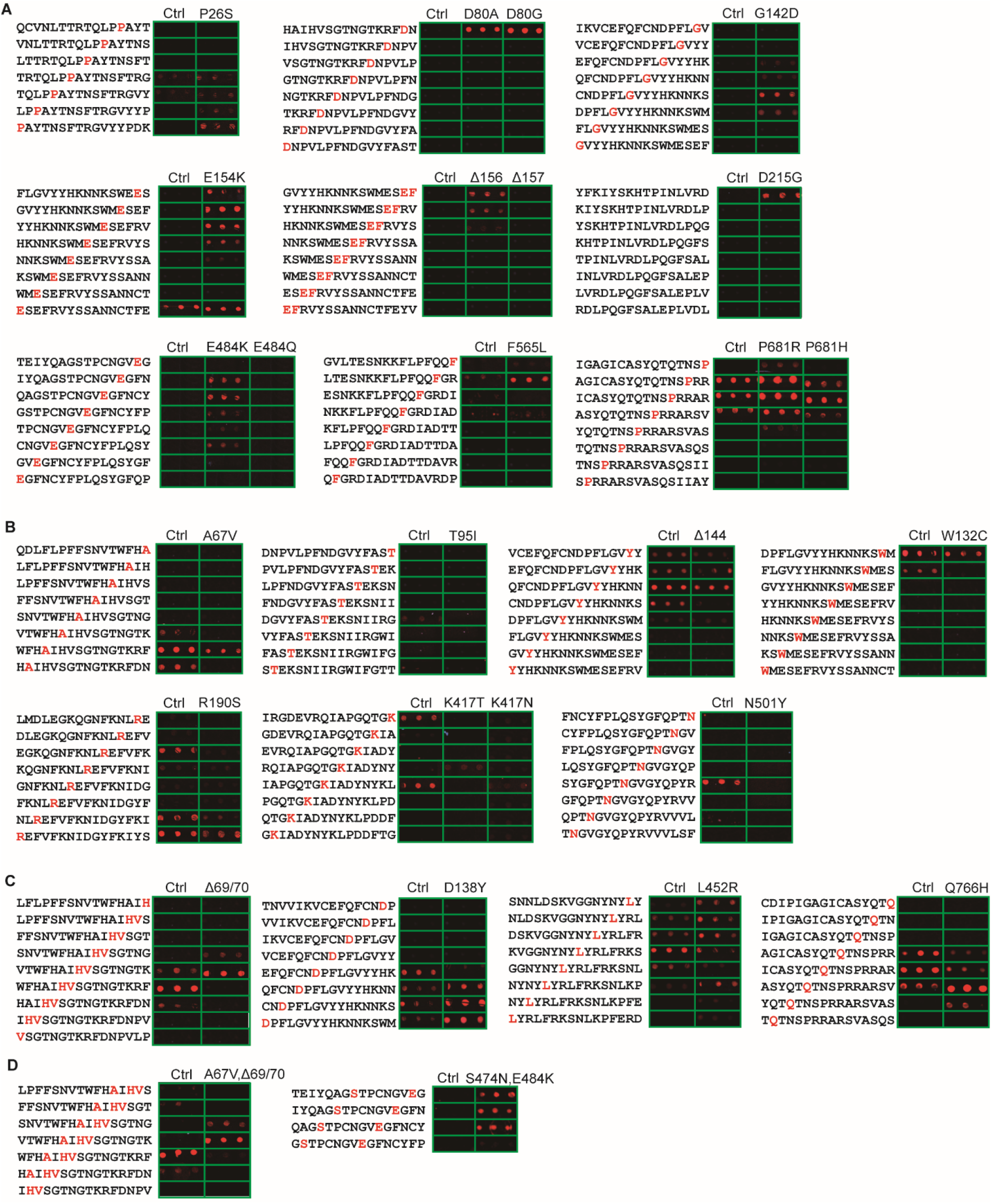
Binding profiles of peptides from SARS-CoV-2 S1 subunits and their mutants to ACE2. Observed intensity increase, decrease, and pattern variation in the array signals were grouped in panels A, B, and C, respectively. (D) Signal change upon multiple amino acid mutations.

The peptide scan has revealed potential additional binding sites of S1 to ACE2. However, we know neither their binding sites on ACE2, nor whether they bind synergistically or competitively to the receptor. In order to assemble the peptides in different combinations and test whether the resulting constructs can recapitulate the multivalent protein-protein interaction to achieve high affinity, we used a peptide-biotin/neutravidin system with four selected peptides. As shown in figure 4A, mixtures of one or multiple biotinylated peptides were mixed with neutravidin. As four biotinylated peptides, same or different, can be displayed on each tetrameric neutravidin, this approach allowed us to test different combinations quickly.

**Figure 4.**
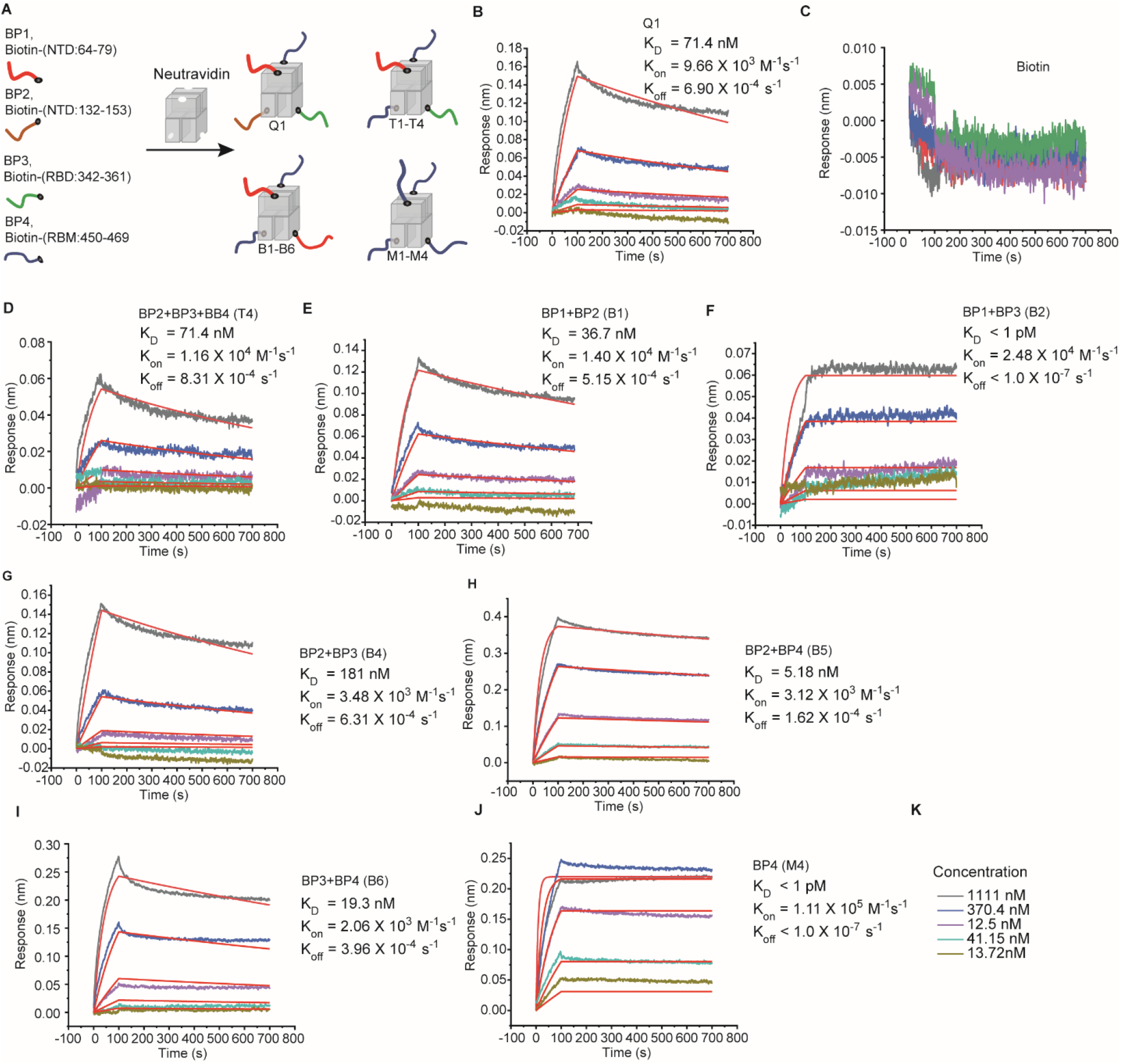
Binding kinetics of biotin-conjugated peptides complexed wih neutravidin to ACE2. (A) The schematic illustration of combinatorially assemblying multiple biotinylated peptides with neutravidin. (B-J) Binding kinetics of biotin-peptide/neutravidin. ACE2 was immobilized on BLI biosensors and measured for binding with various concentrations of the complexes.

To generate the construct randomly displaying four peptides (Q1), the four biotinylated peptides were pre-mixed in equal amounts followed by addition of neutravidin (figure 4A). Its binding to ACE2 was measured using BLI (Bio-Layer Interferometry), showing a K_D_ value of 71.4 nM (figure 4B), remarkably lower than any of the peptides (figure 2). The increased binding affinity is not caused by neutravidin, as no binding can be detected between ACE2 and neutravidin (figure 4C). Among the 4 constructs randomly displaying three peptides, the combination of BP2+BP3+BP4 (T4) exhibits a K_D_ value similar to that of Q1. Remarkably, 5 of the 6 constructs displaying two peptides exhibited strong interactions with ACE2. In particular, BP2+BP4 (B5) exhibited a K_D_ value of 5.2 nM, while the combination of BP1+BP3 (B2) exhibited low pM binding affinity to ACE2. Among the 4 constructs displaying a single peptide, strong binding has been measured only for peptide BP4 (M4). For other combinations, the complexes exhibited strong binding to the reference sensor, so that the affinity cannot be measured reliably.

The dramatic differences between the different constructs remain to be investigated by modelling and structural analysis. The tetrameric neutravidin displays the ligands randomly in a tetrahedron geometry (figure 4A), which can be very different from their arrangement in the folded S1 protein and unfavorable for binding to ACE2. In the future, assembling the peptide fragments in a more spatially defined form, for example, on DNA nanostructures such as DNA origami[16], may represent a more rational approach to designing potent and specific binders. The peptide-biotin/neutravidin method provides a quick screening of different assemblies to identify binders with high affinity comparable to many antibodies. It also demonstrated that the different peptide fragments of S1, upon randomly assembled on a protein template, can recapitulate the protein-protein interaction to achieve high affinity to ACE2.

The interactions between viral molecules and human cells are complex and dynamic. Initiation of infection is often not a simple process of attachment, as many viruses have developed sophisticated mechanisms to evade immune surveillance. For example, the conformational changes involving the different parts of the flexible structure of the spike protein can be associated with the searching of receptors on target cells and escaping immune detection. Therefore, many interactions can take place before the final formation of a stable and specific complex, while these relatively weak and transient interactions cannot be easily accessed by conventional structural biology methods. The fast evolution of viruses also demands a technology, which will allow us to access the influence of the mutations on these interactions quickly. On-demand high-density peptide array synthesis can provide us with a convenient tool to scan these interactions.

## Conclusion

In this work, by using peptide scanning technology, we have identified potential additional receptor binding sites of SARS-CoV and SARS-CoV-2 spike protein subunit 1. Through assembling multiple peptides with a biotin-peptide/neutravidin system, high affinity binders with pM – nM K_D_ have been discovered. In the future, we will synthesize arrays to cover the entire proteomes of corona viruses of different variants, and to probe them not only with human ACE2, but also with other potential human receptors as well as receptors from non-human species. The fast and high throughput synthesis as well as the simple binding-based screening assay will also give us powerful tool to follow and monitor the evolution of the virus. Moreover, this approach will provide new insights into the virus-human cell interactions, not only with regard to the most potent interaction between two specific protein domains, but also to those relative weak and transient in binding. The constructs of assembled and displayed peptide fragments will also be tested, to evaluate their potencies in inhibiting virus attachment and infection.

## Supporting information

peptide sequence summary

## Acknowledgements

We thank Ulrike Hofmann for technical support. This project was supported by the German Bundesministerium für Bildung und Forschung (BMBF) Grant 03Z22E511.

