## Supporting information for "Peptide Scanning of SARS-CoV and SARS-CoV-2 Spike Protein Subunit 1 Reveals Potential Additional Receptor Binding Sites"

### Materials

Acetonitrile, ethanol, Triisopropyl silane (TIPS), Dichloromethane (DCM), 1-[Bis(dimethylamino)methylene]-1H-1,2,3-triazolo[4,5-b] pyridinium 3-oxide hexafluorophosphate (HATU), 1-Hydroxy-7-azabenzotriazol (HOAt), Dithiothreitol (DTT), Dimethyl sulfoxide(DMSO) ACS Gread were purchased from VWR International GmbH (Darmstadt, Germany). Tween 20, N-Methylmorpholine (NMM), N-hydroxysuccinimide (NHS), Chitosan (low molecular weight, 50000 ~ 190000 Da, 75-85% deacetylated, 448869), acetic acid, acetic anhydride, 4-Dimethylaminopyridine (DMAP), HEPES, 5(6)-Carboxyfluorescein, N-hydroxysulfosuccinimide sodium salt (sulfo-NHS) were purchased from Sigma Aldrich. 1-ethyl-3-carbodiimide hydrochloride(EDC), N,N'-Diisopropylcarbodiimide (DIC), D-Biotin, Fmoc-β-Ala-OH, Fmoc-O2Oc-OH, N,N'-Diisopropylethylamine (DIPEA), Dimethylformamide (DMF), Piperidine, Trifluoroacetic acid (TFA), 2-propanol, Fmoc-Rink-Amid Resin, and Fmoc standard amino acids were purchased from IRIS Technology GmbH. DyLight™ 633 NHS Ester, Neutravidin was purchased from Fisher Scientific. Amino functionalized glass slide (Cat.No. 061-1225) was purchased from Tedkon. ACE2 Protein, Human, Recombinant (10108-H08H-B) was purchased from Sino Biological Europe. PD SpinTrap™ G-25 was purchased from GE healthcare.

### Methods

#### SPOT synthesis

The glass slide modification was the same as before, and the peptide array synthesis has some modifications.[1] Biosynthesizer (S/N: 2377-RPC-BS3.2, GeSiM mbH, Germany) was used to automate the synthesis of the high-density peptide array on the glass surface. The glass slide was fixed in the flow cell, which is like a swimming pool to wash glass slide by adding organic solvents, and the organic solvents can be drained like a vacuum filtration system. The procedures were performed cyclically as follows: (1) Fmoc deprotection. 30% piperidine in DMF/Ethanol (volume 1/1) for 30 minutes. After this, the glass slide was washed three times with solvent A (50% DMF/50% Ethanol), then three times with Ethanol. The solvents on the slide were dried by vacuum. (2) Peptide bond formation. Fmoc-amino acid (20 µL,0.25 M in DMSO), HATU (10 µL, 0.5 in DMSO), and NMM (5 µL 3 M in DMF) were freshly mixed. The mixture was printed on the modified glass surface with a volume of 200 pL and a density of 2500 spots per square centimeter at the designed position. The reaction time was 60 mins. After this, the support was washed three times with solvent B (10% acetic acid/45% DMF/45% Ethanol), then three times with solvent A, then three times with Ethanol. The solvents on the support were dried under a vacuum. The coupling process is repeated one more time. (3) Capping of unreacted amino groups. The slide was treated with a fresh mixture of an equal volume of Cap A (10% acetic anhydride in DMF) and Cap B (4% DIPEA in Ethanol) three times, each time 10 mins. After this, the slide was washed three times with solvent A, followed by three times with Ethanol. The solvents on the support are dried by vacuum. After the full-length peptide synthesis finished, the slide was treated with step 1 and step 3, the acetylated the N terminal of the peptides. The glass slide post peptide synthesis is taken out from the flow cell, washed with ethanol, and dried using compressed nitrogen. The glass slide is washed overnight under constant shaking at RT in 2-propanol. On the following day, the cleavage of the side protection groups on amino acids was performed using TFA cocktail: TFA/DTT/Water/TIPS (85/5/5/5) for 2 hours in a sealed bag under constant shaking. The glass slide is washed with ethanol, Milli Q water, and dried using compressed air. The glass slide is cleaned by washing overnight under constant shaking in 2-propanol.

### ACE2 labeling

$NaHCO_3$ was added to the ACE2 solution (50 µL, 0.25 mg/mL in PBS, 5% trehalose, 5% mannitol, 0.01% Tween 80) to reach a final concentration of 100 mM. DyLight™ 633 NHS Ester (0.3 µL, 5 mM in DMSO) was added to the protein solution. The mixture was incubated overnight in the cold room with gentle shaking. Tris (1 M, pH 8.0) was added to a final concentration of 100 mM to quench the reaction for 10 mins, and the free DyLight™ 633 was removed by PD SpinTrap™ G-25.

### Protein Incubation

The glass slide with the chemical library was incubated with ACE2 as described below: A clean glass slide was placed inside the petri dish and the 100µL of 200 nM Dylight 633 labeled ACE2 was gently poured in-between the clean glass slide and the chemical library synthesized glass slide, with the front (the face where the peptide library is synthesized) was facing inwards, such that the protein is sandwiched between the two glass slides. The petri dish was then gently placed in a 4˚C refrigerator overnight. The peptide library synthesized glass slide was gently taken apart from the clean glass slide and immersed in PBS with PBST (0.05% tween 20) buffer for 2 hours. The glass slide was then washed with Milli Q $H_2O$ for 2 mins without shaking, and then gentled dried with kimwipes. The glass slide was then imaged with fluorescence microscopy.

### Peptide synthesis(Table S1)

The peptides were synthesis on Fmoc-Rink-Amid Resin (a scale of 25 µmol) using the Multipep RSI peptide synthesizer. Fmoc standard amino acids, HBTU/Oxyma were dissolved in DMF with a concentration of 0.5M and 45% of NMM in DMF was used as the base. The coupling was performed with 4 eq of Fmoc standard amino acids, 3.8 eq of HBTU/Oxyma, and 12 eq of NMM for 60 mins, and the coupling was performed two times. 20% piperidine in DMF was used for deprotection and 5% acetic anhydride was used for capping. For the normal peptides, the N terminal was acetylated. For biotin-conjugated peptides, the biotin was coupled to the N terminal of the peptide with two repeats of beta-alanine as a linker. After synthesis, the peptides were cleaved from the resin using TFA cocktail (85% TFA (v/v), 5% DTT (w/v), 5% Milli-Q water (v/v) and 5% TIPS (v/v)). The cleaved sample was precipitated and washed with diethyl ether. The sample was dried and dissolved in 200 µL of DMSO. The solution was centrifuged at 13000 rpm for 30 min, and the supernatant was purified using Waters e2695/Waters 2998 equipped with a Luna 100×10mm 5u C18 (S/No:547035-2, Phenomenex Ltd.Germany) column. The column was heated to 50°C during the purification. Waters ACQUITY UPLC Ultra performance/Waters ACQUITY TQ Detector equipped with an analytical C18 column (ACQUITY UPLC BEH C18, bead size 1.7 µm, 50x2.1 mm, Waters, Milford Massachusetts, USA) was used to analyze the mass of peptide synthesized.

### Binding assay by BLI

The Octet RED384 System (FORTEBIO, A SARTORIUS BRAND, GmbH, Germany) equipped with Amine Reactive Second-Generation (AR2G) Biosensors was used to measure the binding affinity. The carboxylic group on the biosensor was activated by 0.1 M EDC and 0.05 M sulfo-NHS for 10 mins, and then reacted with ACE2 (50 µg/mL in PBS/Acetic acid buffer, pH 4.5) overnight in PCR tube, and then quenched with 1 M ethanol amine (1M, pH 8.0) for 10 mins. For the reference sensor, the carboxylic group on the biosensor was activated by 0.1 M EDC and 0.05 M sulfo-NHS for 10 mins, and reacted with 1 M ethanol amine (1M, pH 8.0) for 10 mins. The experiments were performed in HEPES buffer (20 mM HEPES, 150 mM NaCl, 0.05% tween 20, pH 7.4) and the shaking speed was set to 1000 rpm. Various concentration of peptides, or biotin-conjugated peptide/Neutravidin complexes in HEPES buffer was measured for the ACE2 binding on the biosensor. Glycine solution (20 mM, pH 2) was used in between the measurements for biosensor regeneration.

```
SARS-CoV-2      QCVNLTTRTQLPPAYTNSFTRGVYYPDKVFRSSVLHSTQDLFLPFFSNVTWFHAIHVSGT 73
SARS-CoV        RCTTFDDVQAPNYTQHTSSMRGVYYPDEIFRSDTLYLTQDLFLPFYSNVTGFHTIN---- 73
                :*..:        :  .*  *******::***..*: ********:**** **:*:

SARS-CoV-2      NGTKRFDNPVLPFNDGVYFASTEKSNIIRGWIFGTTLDSKTQSLLIVNNATNVVIKVCEF 133
SARS-CoV        ---HTFGNPVIPFKDGIYFAATEKSNVVRGWVFGSTMNNKSQSVIIINNSTNVVIRACNF 130
                   : *.***:**:**:***:*****::***:**:*::.*:**::*:**:*****:.*:*

SARS-CoV-2      QFCNDPFLGVYYHKNNKSWMESEFRVYSSANNCTFEYVSQPFLMDLEGKQGNFKNLREFV 193
SARS-CoV        ELCDNPFFAVSKPMGT----QTHTMIFDNAFNCTFEYISDAFSLDVSEKSGNFKHLREFV 186
                ::*::**:.*    ..    ::.  ::..* ******:*: * :*:.  *.****:*****

SARS-CoV-2      FKNIDGYFKIYSKHTPINLVRDLPQGFSALEPLVDLPIGINITRFQTLLALHRSYLTPGD 253
SARS-CoV        FKNKDGFLYVYKGYQPIDVVRDLPSGFNTLKPIFKLPLGINITNFRAILTAFS------P 240
                *** **:: :*. : **::*****.**.:*:*:..**:*****.*:::*: .

SARS-CoV-2      SSSGWTAGAAAYYVGYLQPRTFLLKYNENGTITDAVDCALDPLSETKCTLKSFTVEKGIY 313
SARS-CoV        AQDIWGTSAAAYFVGYLKPTTFMLKYDENGTITDAVDCSQNPLAELKCSVKSFEIDKGIY 300
                :..  * :.****:****:* **:***:***********: :**:* **::*** ::****

SARS-CoV-2      QTSNFRVQPTESIVRFPNITNLCPFGEVFNATRFASVYAWNRKRISNCVADYSVLYNSAS 373
SARS-CoV        QTSNFRVVPSGDVVRFPNITNLCPFGEVFNATKFPSVYAWERKKISNCVADYSVLYNSTF 360
                ******* *:  .:******************:* *****:**:**************:

SARS-CoV-2      FSTFKCYGVSPTKLNDLCFTNVYADSFVIRGDEVRQIAPGQTGKIADYNYKLPDDFTGCV 433
SARS-CoV        FSTFKCYGVSATKLNDLCFSNVYADSFVVKGDDVRQIAPGQTGVIADYNYKLPDDFMGCV 420
                ********** ********:********::**:********** ************ ***

SARS-CoV-2      IAWNSNNLDSKVGGNYNYLYRLFRKSNLKPFERDISTEIYQAGSTPCNGVEGFNCYFPLQ 493
SARS-CoV        LAWNTRNIDATSTGNYNYKYRYLRHGKLRPFERDISNVPFSPDGKPCTP-PALNCYWPLN 479
                :***:.*:*:.  ***** ** :*:.:*:*******.  :. ...**.   .:***:**:

SARS-CoV-2      SYGFQPTNGVGYQPYRVVVLSFELLHAPATVCGPKKSTNLVKNKCVNFNFNGLTGTGVLT 553
SARS-CoV        DYGFYTTTGIGYQPYRVVVLSFELLNAPATVCGPKLSTDLIKNQCVNFNFNGLTGTGVLT 539
                .***  *.*:***************:********* **:*:**:****************

SARS-CoV-2      ESNKKFLPFQQFGRDIADTTDAVRDPQTLEILDITPCSFGGVSVITPGTNTSNQVAVLYQ 613
SARS-CoV        PSSKRFQPFQQFGRDVSDFTDSVRDPKTSEILDISPCAFGGVSVITPGTNASSEVAVLYQ 599
                 *.*:* ********::* **:****:* *****:**:************:*.:******

SARS-CoV-2      DVNCTEVPVAIHADQLTPTWRVYSTGSNVFQTRAGCLIGAEHVNNSYECDIPIGAGICAS 673
SARS-CoV        DVNCTDVSTAIHADQLTPAWRIYSTGNNVFQTQAGCLIGAEHVDTSYECDIPIGAGICAS 659
                *****:* .**********:**:****.*****:**********:.***************

SARS-CoV-2      YQTQTNSPRRARSVASQSIIAY 695
SARS-CoV        YHTVSLLRST------------ 669
                *:*  :
```

**Figure S1. Sequence alignment of SARS-CoV-2 and SARS-CoV S1 subunits.** The peptides with positive signals were highlighted with yellow color. The RBM motifs were indicated as red.

**Table S1. Binding kinetics of peptides from SARS-CoV-2 to ACE2**

| SARS-CoV-2 | $K_D$ (M) | $K_D$ Error | $k_{on}$ (1/Ms) | $k_{on}$ Error | $k_{off}$ (1/s) | $k_{off}$ Error |
|---|---|---|---|---|---|---|
| NTD:64-79 | 2.31E-05 | 2.09E-06 | 5.32E+03 | 3.91E+02 | 1.23E-01 | 6.51E-03 |
| NTD:132-153 | 1.22E-05 | 3.99E-07 | 5.12E+03 | 1.38E+02 | 6.24E-02 | 1.17E-03 |
| NTD:188-219 | 5.20E-05 | 1.32E-05 | 1.04E+03 | 2.60E+02 | 5.41E-02 | 2.09E-03 |
| RBD:342-361 | 5.29E-07 | 2.65E-08 | 1.71E+04 | 7.82E+02 | 9.05E-03 | 1.88E-04 |
| RBM:450-469 | 4.00E-04 | 7.50E-05 | 1.62E+03 | 2.70E+02 | 6.49E-01 | 5.63E-02 |

**Table S2. Binding kinetic of peptides from SARS-CoV to ACE2**

| SARS-CoV | $K_D$ (M) | $K_D$ Error | $k_{on}$ (1/Ms) | $k_{on}$ Error | $k_{off}$ (1/s) | $k_{off}$ Error |
|---|---|---|---|---|---|---|
| **RBD: 328-349** | 3.66E-05 | 2.35E-06 | 2.17E+03 | 1.22E+02 | 7.94E-02 | 2.42E-03 |
| **RBD: 358-375** | 2.01E-06 | 3.46E-08 | 6.27E+02 | 8.04E+00 | 1.26E-03 | 1.45E-05 |
| **RBM: 426-453** | 1.88E-05 | 2.07E-06 | 3.95E+03 | 3.62E+02 | 7.41E-02 | 4.55E-03 |

**Table S3. Binding kinetic of biotin-conjugated peptides complexed with neutravidin to ACE2**

| Combinations | $K_D$ (M) | $K_D$ Error | $k_{on}$ (1/Ms) | $k_{on}$ Error | $k_{off}$ (1/s) | $k_{off}$ Error |
|---|---|---|---|---|---|---|
| **BP4** | <1.0E-12 | 8.79E-11 | 1.11E+05 | 8.46E+02 | <1.0E-07 | |
| **BP1 & BP2** | 3.67E-08 | 7.73E-10 | 1.40E+04 | 1.49E+02 | 5.15E-04 | 9.37E-06 |
| **BP1 & BP3** | <1.0E-12 | 6.99E-10 | 2.48E+04 | 3.47E+02 | <1.0E-07 | |
| **BP2 & BP3** | 1.81E-07 | 9.85E-09 | 3.48E+03 | 1.76E+02 | 6.31E-04 | 1.24E-05 |
| **BP2 & BP4** | 5.18E-09 | 1.03E-10 | 3.12E+04 | 7.50E+01 | 1.62E-04 | 3.19E-06 |
| **BP3 & BP4** | 1.93E-08 | 3.86E-10 | 2.06E+04 | 1.36E+02 | 3.96E-04 | 7.50E-06 |
| **BP2 & BP3 & BP4** | 7.14E-08 | 1.73E-09 | 1.16E+04 | 2.06E+02 | 8.31E-04 | 1.36E-05 |
| **BP1 & BP2 & BP3 & BP4** | 7.14E-08 | 1.27E-09 | 9.66E+03 | 1.25E+02 | 6.90E-04 | 8.45E-06 |

**Table S4. Summary of synthesized peptides.**

| Peptide sequences | Molecular weight | Observed Molecular weight | Source | Location | Array Location |
|---|---|---|---|---|---|
| WFHAIHVSGTNGTKRF | 1899.12 | 1898.50 | SARS-CoV-CoV-2 | NTD:64-79 | B9 |
| SLLIVNNATNVVIKVC | 1741.11 | Can’t dissolve | SARS-CoV-CoV-2 | NTD:116-131 | D1 |
| EFQFCNDPFLGVYYHKNNKSWM | 2809.14 | 2080.07 | SARS-CoV-CoV-2 | NTD:132-153 | D9-D12 |
| NLREFVFKNIDGYFKIYSKHTP | 2758.13 | 2756.89 | SARS-CoV-CoV-2 | NTD:188-219 | F3-F6 |
| DCALDPLSETKCTLKSFT | 2013.3 | 2012.24 | SARS-CoV-CoV-2 | NTD:290-307 | I3-I4 |
| FNATRFASVYAWNRKRISNC | 2445.76 | 2444.82 | SARS-CoV-CoV-2 | RBD:342-361 | J12-J14 |
| EVRQIAPGQTGKIADY | 1786.98 | 1786.16 | SARS-CoV-CoV-2 | RBD:406-421 | L10 |
| NLDSKVGGNYNYLYRLFRKS | 2448.73 | 2447.53 | SARS-CoV-CoV-2 | RBM:450-469 | M10-M12 |
| SFELLHAPATVCGPKKST | 1927.23 | 1926.57 | SARS-CoV-CoV-2 | RBD:514-531 | O13-O14 |
| VCGPKKSTNLVKNKCVNFNF | 2281.7 | 2281.38 | SARS-CoV-CoV-2 | RBD:524-543 | P1-P3 |
| DAFSLDVSEKSGNFKHLREFVFKNKD | 3099.41 | 3100.12 | SARS-CoV | NTD:166-191 | E7-E12 |
| VRDLPSGFNTLKPIFK | 1873.2 | 1872.27 | SARS-CoV | NTD:206-221 | F10 |
| IWGTSAAAYFVGYLKPTT | 1987.26 | 1987.40 | SARS-CoV | NTD:244-261 | G12-G13 |
| VDCSQNPLAELKCSVKSF | 2009.31 | 2008.34 | SARS-CoV | NTD:276-293 | H11-H12 |
| VFNATKFPSVYAWERKKISNCV | 2629.04 | 2628.02 | SARS-CoV | RBD:328-349 | J3-J6 |
| STFFSTFKCYGVSATKLN | 2042.31 | 2042.43 | SARS-CoV | RBD:358-375 | K1-K2 |
| RNIDATSTGNYNYKYRYLRHGKLRPFER | 3530.91 | 3529.34 | SARS-CoV | RBD:426-453 | M1-M7 |
| SFELLNAPATVCGPKLSTDLIKNQCV | 2803.26 | 2802.98 | SARS-CoV | RBD-SD1:500-525 | O4-O9 |
| NGLTGTGVLTPSSKRF | 1675.88 | 1675.33 | SARS-CoV | SD1:530-545 | P2 |
| LTPSSKRFQPFQQFGR | 1965.22 | 1964.40 | SARS-CoV | SD1:538-553 | P6 |
| VSTAIHADQLTPAWRIYSTG | 2228.46 | 2227.4 | SARS-CoV | SD2:606-625 | R6-R8 |
| Biotin-bA[1]-bA-WFHAIHVSGTNGTKRF | 2225.54 | 2224.45 | SARS-CoV-CoV-2 | NTD:64-79 | B9 |
| Biotin-bA-bA-EFQFCNDPFLGVYYHKNNKSWM | 3135.56 | 3134.51 | SARS-CoV-CoV-2 | NTD:132-153 | D9-D12 |
| Biotin-bA-bA-FNATRFASVYAWNRKRISNC | 2772.18 | 2770.96 | SARS-CoV-CoV-2 | NTD:342-361 | J12-J14 |
| Biotin-bA-bA-NLDSKVGGNYNYLYRLFRKS | 2775.15 | 2774.02 | SARS-CoV-CoV-2 | NTD:450-469 | M10-M12 |

bA, beta Alanine

1. Lin, W., et al., *Controlling Surface Wettability for Automated In Situ Array Synthesis and Direct Bioscreening.* Advanced Materials. **n/a**(n/a): p. 2102349.
